# T2 heterogeneity provides a sensitive measure of early tumor response to radiotherapy

**DOI:** 10.1101/2020.04.21.053736

**Authors:** Michal R. Tomaszewski, William Dominguez-Viqueira, Antonio Ortiz, Yu Shi, James R. Costello, Heiko Enderling, Stephen A. Rosenberg, Robert J. Gillies

**Affiliations:** Department of Cancer Physiology, H. Lee Moffitt Cancer Center and Research Institute; Small Imaging Laboratory Core Facility, H. Lee Moffitt Cancer Center and Research Institute; Analytical; Microscopy Core Facility, H. Lee Moffitt Cancer Center and Research Institute; Department of Radiology, ShengJing Hospital of China Medical University; Department of Radiology, H. Lee Moffitt Cancer Center and Research Institute; Department of Integrated Mathematical Oncology, H. Lee Moffitt Cancer Center and Research Institute; Department of Radiation Oncology, H. Lee Moffitt Cancer Center and Research Institute

**Keywords:** MRI, tumor heterogeneity, radiotherapy, pancreatic cancer

## Abstract

**Purpose:** External beam radiotherapy (XRT) is a widely used cancer treatment, yet responses vary dramatically between patients. These differences are not accounted for in clinical practice, in part due to a lack of sensitive biomarkers of early response. In this work, we test the hypothesis that quantification of intratumor heterogeneity is a sensitive and robust biomarker of early response to XRT. A novel Magnetic Resonance Imaging (MRI) approach is proposed, utilizing histogram analysis of clinically-used T2 relaxation measurements to assess early changes in the tumor heterogeneity following irradiation in murine models of pancreatic cancer, indicative of radiotherapy response.

**Methods and Materials:** Dynamic Magnetic Resonance T2 relaxation imaging was performed every 72h following 10 Gy dose XRT in two murine models of pancreatic cancer. Proposed biomarker of radiotherapy response was compared with tumor growth kinetics, and biological validation was performed through quantitative histology analysis.

**Results:** Quantification of tumor T2 interquartile range (IQR) as a measure of histogram width showed excellent sensitivity for detection of XRT-induced tumor changes as early as 72h after treatment, outperforming whole tumor T2 and Diffusion weighted MRI metrics. This response was observed both in quantitative T2 maps and in T2-weighted images that are routine in clinical practice. Histological comparison revealed the T2 IQR provides a measure of spatial heterogeneity in tumor cell density, related to radiation-induced necrosis. The early IQR changes were found to presage subsequent tumor volume changes in two distinct pancreatic models, suggesting promise for treatment response prediction. The metric showed excellent test-retest robustness.

**Conclusions:** Our preclinical findings indicate that spatial heterogeneity analysis of T2 MRI can provide a sensitive and readily translatable method for early radiotherapy response assessment in pancreatic cancer. We propose that this will be useful in adaptive radiotherapy, specifically in MRI-guided treatment paradigms.

## Introduction

Rapid growth and continuous evolution of tumors in response to changing microenvironmental pressures leads to significant intratumor heterogeneity in cell distribution, micro-environmental characteristics, and genomics (1). The importance of accurate measurement of tumor heterogeneity for understanding disease progression and treatment response is becoming increasingly appreciated. However, sensitive and validated techniques that exploit the information available from quantification of inhomogeneous internal tumor structure are still lacking in clinical practice (2,3). In particular, measurement of tumor response to chemo- and radio-therapeutic interventions, strongly affecting the internal structure and heterogeneity of the lesion, is of paramount importance for optimization of treatment and improvements in patient outcomes.

External beam radiotherapy (XRT) is widely used in treatment of multiple cancer types with the aim of local control of the rapidly dividing tumor cells through delivery of ionizing radiation and subsequent induction of DNA damage (4). This simplified mechanism of action is complicated due to multiple factors that are often heterogeneously expressed within the same tumor. This intratumoral heterogeneity can lead to differences in response that are not accounted for in clinical practice, wherein the great majority of patients receive standardized XRT routines. In this work, we propose a novel, clinically applicable Magnetic Resonance Imaging (MRI) approach to assess early changes in tumor heterogeneity in response to irradiation, which may provide information to adapt treatment and improve outcomes.

Magnetic Resonance methods have been used for radiotherapy response measurement. Diffusion and T2 imaging serve as a measure of structural changes (5,6) due to cell death, while functional MR techniques can be used to document acute vascular damage. These include (7) Dynamic Contrast Enhanced (DCE) (8), Oxygen-Enhanced (OE) (9,10) and intrinsic susceptibility (IS) (11) MRI. However, these methods are rarely used for clinical decision support, in part reflecting intrinsic poor sensitivity of MRI as well as relative sophistication of the protocols and long acquisition times involved. Novel analytic approaches may address some of these challenges. It has been postulated that metrics describing the internal structure and heterogeneity within tumors may provide more information about the disease than traditional whole-tumor metrics (12,13). Characterization of tumor heterogeneity with MR imaging has been attempted previously using texture analysis (12,14,15), clustering methods (16–19), and other structural descriptors (20). However, such efforts often lack biological interpretation and clear clinical relevance.

Pancreatic cancer may benefit particularly from the advances in MR-based therapy response assessment. Locally advanced Pancreatic Ductal Adenocarcinoma (PDAC) has a poor 5-year survival rate of around 9%. (21). Approximately 1/3 of patients will die of local progression (22). Evaluating therapeutic response of local disease as early as possible is therefore crucial, as maximization of the therapeutic impact to the primary site is critical. Increasingly, data are showing the promise of radiotherapy (23) for treatment of locally advanced PDAC and, in particular, MR-guided stereotactic body radiation (24). The advent of the MR guided radiotherapy (MRgRT) systems (25), supplies a dynamic imaging record of tumor changes with every radiation fraction, providing an opportunity for a dramatic increase in the role of MR in XRT response detection. To date, these valuable image data have been used purely for tissue segmentation and beam positioning. However, we propose that these image data could also be used to assess intermediate responses and modulate the next doses accordingly. To maximize translation potential for all MRgRT systems (26,27), robust sequences suitable for low field scanners should be considered. We believe that the method described below has the potential to address this unmet need, utilizing clinically acquired anatomic MRI data to provide a sensitive measure of response.

Herein, we show in a murine model of PDAC that quantification of T2 histogram interquartile range (IQR) reflects tumor heterogeneity and shows sensitivity to early XRT response that are superior to standard whole-tumor T2 and apparent diffusion coefficient (ADC)-based metrics previously used for detection of radiation-induced necrosis. Importantly, histological insight into the source of the observed changes is provided. In addition, results highlight that this approach can be used in conjunction with simple T2 weighted imaging, routinely used for anatomical MRI. The findings are validated in a separate tumor model, and proof-of-concept clinical results are presented.

## Methods

### Cells and Tumors

All procedures were approved by the Institutional Animal Care and Use Committee (IACUC), at our institution under the protocol 4778. Mouse pancreatic adenocarcinoma Panc02 cells were a kind gift from Emmanuel Zervos (East Carolina University, Greenville, NC). Human pancreatic adenocarcinoma BXPC3 cells were obtained from American Type Culture Collection, ATCC (Rockville, MD). These were cultured in RPMI 1640 cell culture media supplemented with 10% Fetal Bovine Serum and passaged weekly. Both cell lines were mycoplasma free and have been authenticated using short tandem repeat (STR) DNA profiling. At the time of inoculation, cells suspended in 0.1 mL 50:50 PBS and Matrigel (Corning, NY), and implanted subcutaneously into upper hind legs of mice. 2×10^6^ Panc02 were injected into BL/6 mice (n=29), and 5×10^6^ BXPC3 cells were injected into immunodeficient NOD SCID gamma (NSG) mice (n=12). Mice were obtained from our breeding colonies from breeding pairs that were established and are refreshed every 6 months from Jackson Labs, JAX (Bar Harbor, ME). When the tumors reached approximately 8 mm linear dimension as measured by calipers, mice were randomly allocated to treated and control groups.

### MR Imaging

All animals underwent MRI for tumor size measurement 3 days before irradiation, followed by full MRI protocol as described below, first 2h prior to irradiation (sham irradiation for control animals) and every 3 days thereafter. Following the last imaging (day 9 and 12 respectively for Panc02 and BXPC3 cohorts), the mice were sacrificed and tumors excised. One Panc02 tumor bearing animal had to be euthanized prior to the last time-point due to tumor ulceration. 20 Gy treated mice were only imaged on day 0 and day 3 after irradiation to capture early changes. Details of MR imaging are presented in Supplementary Methods.

### XRT

Mice were anaesthetized using intraperitoneal injection of ketamine (10mg/kg) and xylazine (1mg/kg) mixture and placed in the irradiator chamber (XRAD 320, Precision X-Ray, CT) with lead shielding covering the entire body, only exposing the tumor on the leg. X-Ray dose was delivered at a rate of 152 cGy/min (320 kV, 12.5A, 1.3 min, 6.6 min and 13.2 min for 2 Gy, 10 Gy and 20 Gy doses, respectively).

N=11 Panc02 tumors were irradiated with 10 Gy dose in single fraction and smaller cohorts received a single 20 Gy dose (n=6) or 5 daily doses of 2Gy (n=6) to evaluate the effect of dose and fractionation, while n=6 received sham irradiation. For validation, mice bearing BXPC3 tumors received single 10 Gy irradiation dose (n=8) or sham irradiation (n=4). For test-retest analysis, n=4 Panc02 and n=4 BXPC3 tumor bearing mice were imaged twice with the standard MRI protocol. Between the two scans the animal was removed from the holder, checked visually for approximately 10s and placed back in the holder, meaning that the tumor positioning was slightly different. Coil tuning and matching, shimming and reference power calculation were performed independently for the test and retest scans, and regions of interest were drawn separately from the corresponding anatomical T2 weighted scans.

Histology. Following excision, tumors were marked with tissue dyes (Cancer Diagnostics Inc., NC, one green line marking the top, one black line marking the left side) to preserve orientation. Tumors were fixed (10% Neutral Buffered Formalin, 24h) and cut parallel to the imaging plane into 4 mm thick slices, which were paraffin embedded, with 5um sections cut and stained with Hematoxylin and Eosin (H&E) to reveal tissue structure and viability. H&E stained sections were segmented and analyzed to quantify local nuclear density distribution, as detailed in Supplementary Methods.

### Image Analysis

All MRI image analyses were performed in MATLAB 2018b (Mathworks, Natick, MA) using custom written code. Tumors were outlined manually in the high-resolution MRI T2 weighted images, with the same masks used for analysis of T2 and ADC maps. The histogram metrics (mean, median, standard deviation, kurtosis, skewness, interquartile range (IQR)) were computed for the voxel populations inside the tumor masks using built-in MATLAB functions (*mean(), median(), std(), kurtosis(), skewness(), iqr()* respectively). Representative slices of qT2 maps were matched with the corresponding H&E sections based on the slice position and histology dye marks, enabling identification and segmentation of necrotic and viable areas in the MRI images. T2 image value statistics were then compared in these regions. Raw single echo images acquired for quantification of the T2 relaxation time were also analyzed as a simulation of T2 weighted images, as commonly acquired in clinical practice. The scans were compared at each echo time (32 echoes 7-224 ms TE in 7 ms intervals), with median and IQR computed as for T2 maps, and the changes of the metrics compared before and after XRT to assess the sensitivity to irradiation response as a function of the echo time.

### Statistical analysis

All errors are quoted as the SEM unless otherwise stated. All statistical analyses were performed in MATLAB 2018b and MS Excel 2010 (Microsoft, Redmond, WA). Paired two-tailed t-test compared metrics at different time-points, unpaired two-tailed t-test compared metrics between different cohorts. Pearson rank test was performed to assess significance of correlation between parameter changes and treatment response. 1-to-1 Interclass correlation coefficient (28) was used to assess repeatability in test-retest experiments. P < 0.05 was considered statistically significant.

### Human image analysis

Proof-of-conceot quantification of histogram Interquartile range as a surrogate measure of tumor heterogeneity was performed on T2 weighted MRI scans from two PDAC patients. The Institutional Review Board at our institution approved (IRB #17837) and waived the informed consent requirement for retrospective analysis in this study. Axial T2 weighted MRI scans of two advanced PDAC patients was acquired at our institution on a 1.5T MRI scanner (GE Medical Systems, Waukesha, WI) using a Fast Spin Echo sequence. The lesions were segmented in each slice by an experienced radiologist in HealthMyne software (HealthMyne Inc, Madison, WI), and transferred into MATLAB, where quantification was performed.

## Results

### Interquartile range of T2 shows excellent sensitivity to early radiotherapy response

The Panc02 tumors showed a rapid volume increase, as seen in the 3 days prior to irradiation, that was significantly slowed within the first 3 days after the 10 Gy dose to the tumor (1.22±0.07 vs 1.65±0.13 3-day volume increase ratio, p<0.001, **Figure 1A**). This slowdown persisted for another 3 days (1.11±0.04 3 day volume increase ratio, p<0.001 to pre-XRT) with the tumors resuming significantly faster growth between days 6 and 9 after irradiation. There was a 33 ± 6% volume increase between days 6 and 9, compared to the 11 ± 4% volume increase between days 3 and 6 (p=0.008).

**Fig. 1:**
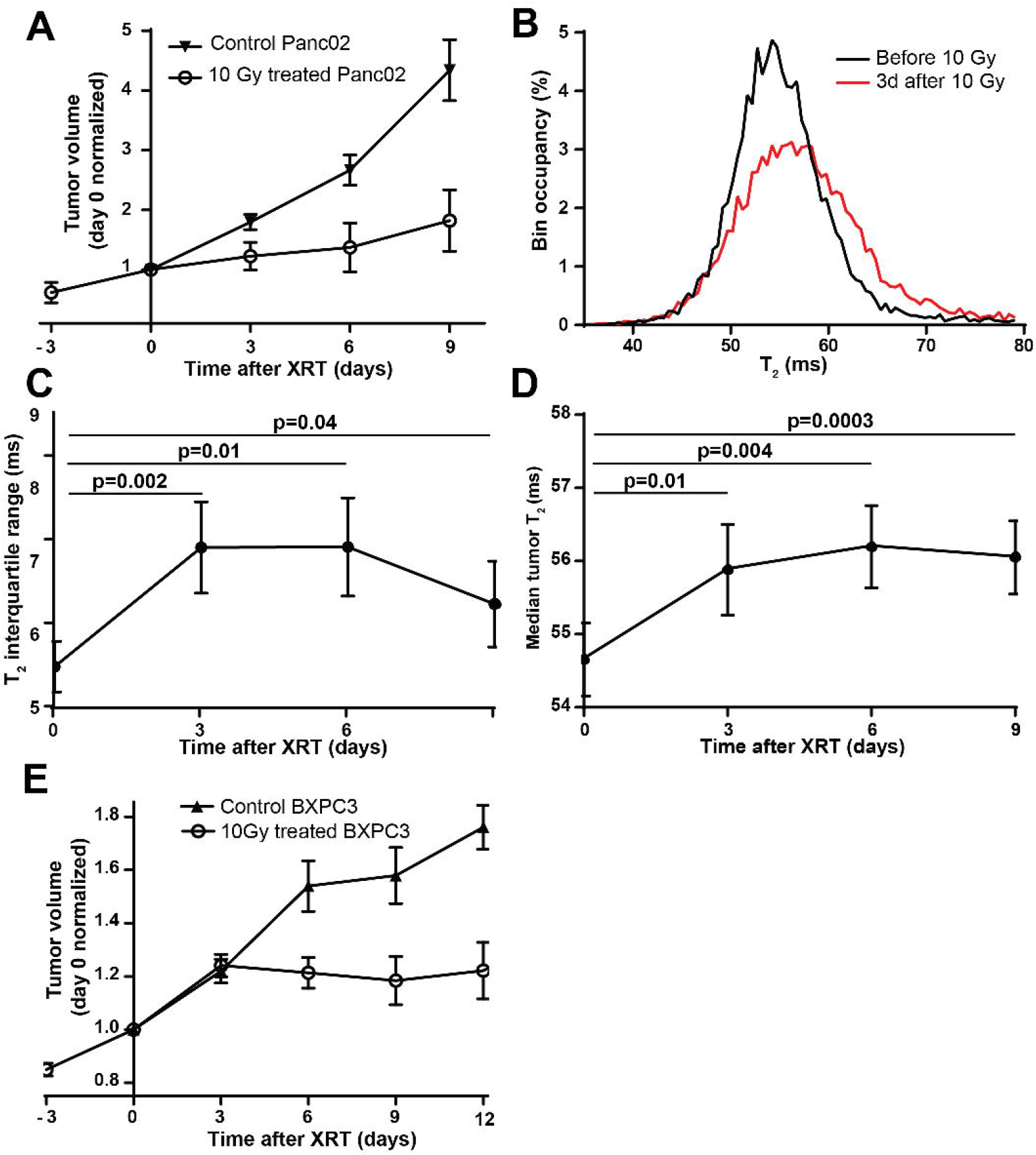
Radiotherapy causes dramatic changes in tumor T2 maps. (A) Irradiated Panc02 tumors slow down their volume increase, as measured by MRI, with faster growth returning 6 days after irradiation. (B) Quantification of the histogram of T2 relaxation times in representative tumor voxels show a change in the histogram after 3 days (40 bins, normalized to Area Under Curve=1). Quantification of the histogram metrics shows that these changes are best described by the Interquartile Range and median. Plotted every 3 days after irradiation, (C) a significant early increase in T2 interquartile range is observed, while the T2 median (D) shows modest early changes. 10 Gy treated BXPC3 tumors also show growth slowdown (E) compared to control, with some growth returning after 9 days, n=11 treated and n=6 untreated Panc02 tumors (A-D), n=8 treated and n=4 untreated BXPC3 tumors (E), error bars denote standard errors of the mean. Volumes were normalized to value at day 0 and averaged to display the growth trend.

Comparison of the T2 histograms suggests complex changes in value distribution accompany the growth slow-down as early as 72h post-irradiation. A representative histogram pair (**Figure 1B**), shows a clear broadening of value distribution coupled with a slight shift to higher values. Quantification of the relevant metrics confirms these observations. We observed a strong, highly significant increase in the width of the histogram, expressed as the interquartile range (IQR, 6.9±0.5ms vs. 5.4±0.3ms, p<0.001, n=11, **Figure 1C**). A small increase in median T2 (55.8±0.6ms vs. 54.6±0.5ms, p=0.01, n=11, **Figure 1D**), and a related slight but significant increase in the mean (58.3±0.9ms vs. 56.0±0.7ms, p=0.04, n=11) described the shift. The effects of irradiation on other standard histogram metrics were not significant (See **Supplementary Table 1**). IQR showed different temporal evolution profiles than the median, suggesting a different informational content between these metrics. Consistent with the tumor growth resuming after day 6 (**Figure 1A**), marking the return of tumor cells toward a normal proliferation pattern, the IQR also dropped back towards baseline at this point, whereas the change in the median between days 6 and 9 was not significant. Non-irradiated animals showed no significant changes in the imaging metrics (**Supplementary Table 2**).

Diffusion weighted Imaging, routinely used for detection of macroscopic tumor necrosis, performed worse in the above setting, with the Apparent Diffusion Coefficient (ADC) showing weaker, nonsignificant trends in any of the considered metrics following 10 Gy irradiation (eg. Median: 559±24 10^-6^ vs. 513±29 10^-6^ mm^2^/s, p=0.21, IQR: 232±26 10^-6^ vs. l85±l9 10^-6^ mm^2^/s, p=0.11, 3d after vs. before 10 Gy XRT, n=11). Escalation of the dose to 20 Gy, beyond standard clinical practice, was used to verify that the small measured change was related to the relatively poor sensitivity of the method, producing a significant change in median ADC as expected (619±19 vs. 521±20 10^-6^ mm^2^/s 3d after vs. before 20Gy XRT, p=0.019 n=6, see **Supplementary Figure S1**), while still underperforming T2 in the same cohort (median 58.2±0.8 vs. 55.7±0.9 ms, p=0.0008, n=6).

Another, smaller cohort of tumor-bearing mice was subject to a traditionally fractionated 2Gy x 5 days radiotherapy regime in order to further test the utility of IQR for early response measurement, as observed in the single 10 Gy dose cohort. The results (**Supplementary Figure S2**) show an even more strikingly superior performance of IQR vis-a-vis median T2. At day 3, after 6 Gy of the 10 Gy total dose, when no significant change in tumor growth dynamics were yet observed (1.62±0.06 vs. 1.84±0.11 3 day volume increase ratios, p=0.24), the median T2 remained unchanged, yet the IQR showed a clear, nearly significant increase between these time-points (5.8±0.4 vs. 5.2±0.5 ms, p=0.06), again indicating the utility of the metric for high-sensitivity detection of early response.

An alternative tumor model, a less aggressive, human PDAC cell line, BXPC3, was used to test these findings. As expected, a clearer growth arrest was observed (**Figure 1E**) in these slower growing, well-differentiated tumors, with total growth arrest appearing after 6 days (volume ratio over 3 days 0.97±0.03, vs. 1.18±0.05 before treatment, p=0.01), and persisting until day 9 (1.02±0.04 volume ratio). Observing the T2 changes, as for Panc02, the IQR showed significant increase directly after XRT unlike any of the other histogram metrics (**Supplementary Table 3**). Hence, two different models with different growth characteristic both showed the same pattern of change in histogram metrics. No significant changes in histogram metrics were observed in non-irradiated BXPC3 tumors (**Supplementary Table 4**).

### Early Interquartile range changes presage tumor volume regrowth

It was hypothesized that these early change in T2 IQR in response to irradiation, as observed in both tumor types, may be indicative and provide insight into the longer-term tumor size response. Consistent with this hypothesis, the ratio of IQR increase 3 days after 10 Gy single dose irradiation was inversely correlated with the tumor growth between day 6 and 9 in Panc02 tumors, when re-growth was observed (Pearson r=-0.67, p=0.035, **Figure 2A**). These data show that a stronger early IQR response presages a slower tumor re-growth, suggesting that the IQR changes may have predictive value for tumor XRT response.

**Fig. 2:**
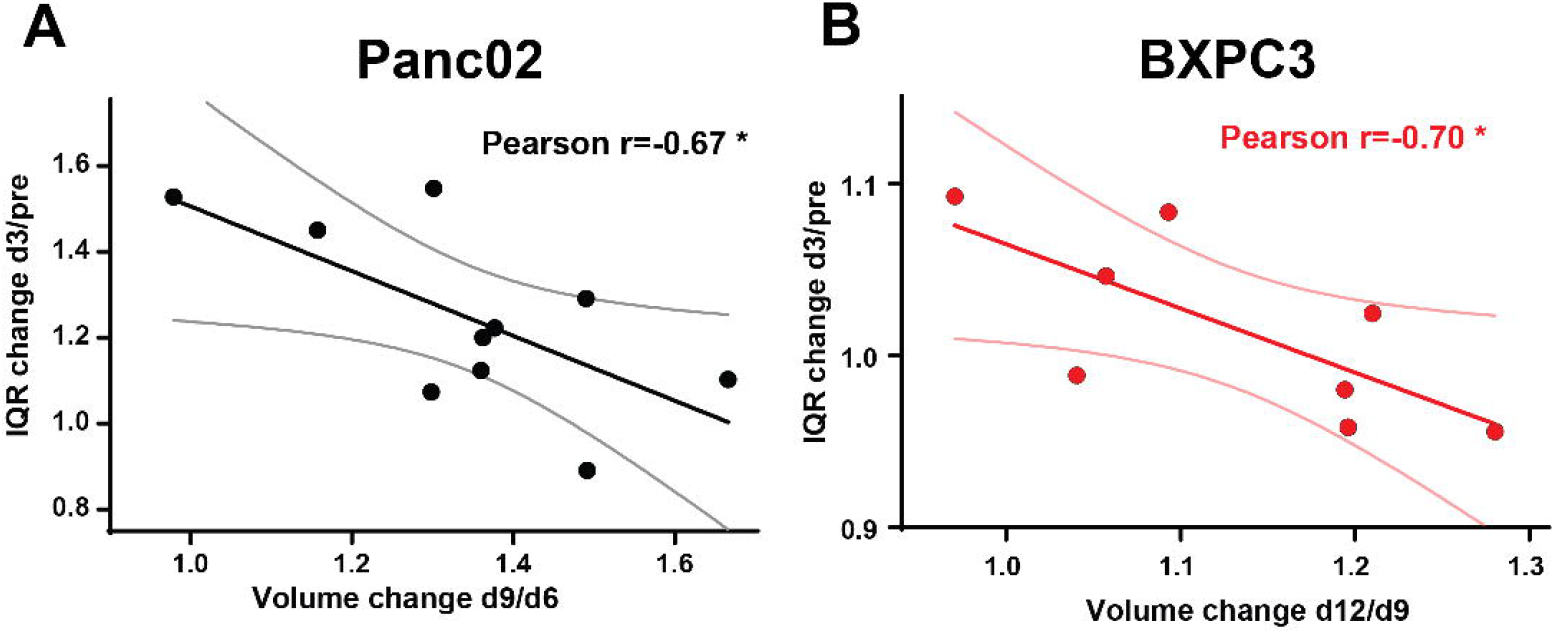
Early changes in T2 interquartile range correlate to later tumor growth after radiotherapy. A significant negative correlation was observed between the interquartile range ratio 72h after/before 10 Gy irradiation, and volume re-growth defined as the volume ratio 9 days/6 days after irradiation (Panc02 tumors, A) or 12 days/9 days after irradiation (BXPC3, B). The time-points were chosen based on when tumors return to fast growth (simulating recurrence). N=10 Panc02 and n=8 BXPC3 tumors, line of best fit and 95% confidence intervals plotted, Pearson r correlation coefficients quoted.

Strengthening this observation, the correlation between IQR change and volume re-growth was also observed in BXPC3 tumors. In this case, as mentioned above, a significant acceleration of growth was observed 9-12d after irradiation. The volume ratio between these time-points was therefore compared to the 3d IQR increase, revealing a significant negative correlation (Pearson r=-0.70, p=0.036, **Figure 2B**). Importantly, the early size change itself did not significantly correlate with the regrowth in either tumor type (Pearson r=-0.20, p=0.56 and r=0.14,p=0.72).

### Tumor necrosis and structural heterogeneity drives the T2 IQR metric

We then investigated the biological interpretation of the results, in order to better understand the reason behind the high sensitivity of structural heterogeneity as a measure of radiotherapy response.

The two tumor types displayed clear differences in baseline imaging metrics. The bright, heterogeneous appearance of the BXPC3 tumors (**Figure 3A**) in T2 maps is apparent in the representative T2 histograms (**Figure 3B**) and confirmed in a significantly higher T2 IQR (12.6±0.6ms vs 5.5±0.3ms, **Figure 3C**) as well as higher median (58.4±0.6ms vs 54.6±0.5ms, **Figure 3D**) in the entire cohorts at day 0 compared to Panc02. The change in IQR following 10 Gy irradiation was significantly higher in the Panc02 tumors (1.25±0.06 vs 1.10±0.03 ratio 3d/pre, **Figure 3E**).

**Fig. 3:**
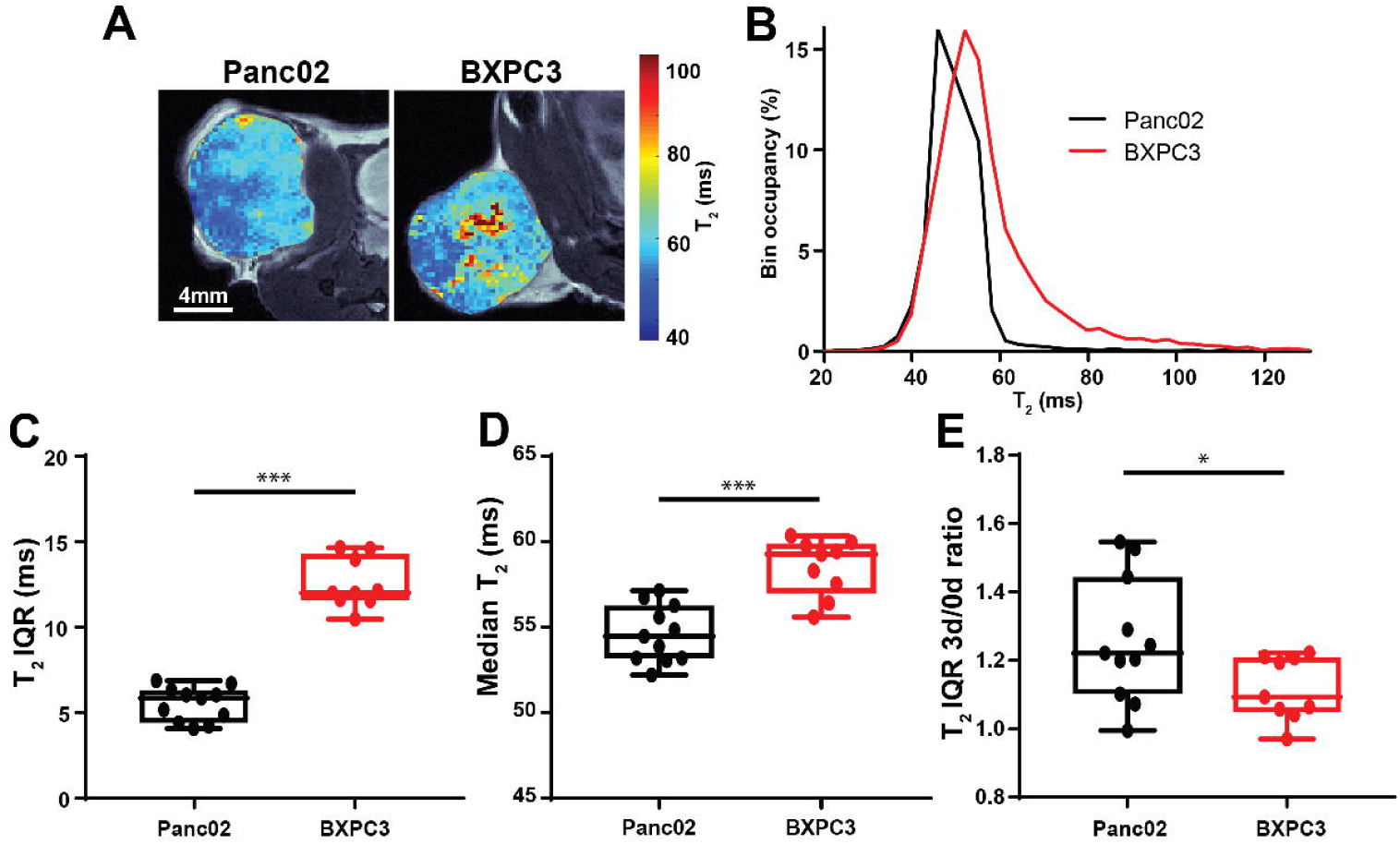
Panc02 and BXPC3 tumors show contrasting T2 metrics. Representative Panc02 (A,left) tumor shows more homogenous distribution and lower values of T2 relaxation time to BXPC3 tumor (A,right). As apparent from representative T2 value histograms (B), quantification of the interquartile range (IQR,C) and median (IQR,D) of T2 confirms these observations. A significantly stronger IQR change after 10 Gy irradiation (3 days/pre XRT ratio), is observed in Panc02 tumors (E). Box between 25th and 75th percentile, line at median, n=11 Panc02, n=8 BXPC3. *:p<0.05, ***: p<0.001

Comparison of the histological quantification of average nuclear densities in images down-sampled to match MR resolution of a representative subset of tumors reflected the observed differences in MR parameters. Clearly visible in the representative images (**Figure 4A**) and corresponding histograms (**Figure 4B**), the BXPC3 tumors exhibited significantly higher IQR of nuclear density (2.4±0.3% vs. 1.12±0.07%, p=0.01, **Figure 4D**) and lower median (8.41±0.13% vs. 10.28±0.07%, p=0.0001, **Figure 4C**) than Panc02 tumors. These findings were consistent with the significantly elevated T2 IQR and T2 median in BXPC3 vis-à-vis Panc02 tumors.

**Fig. 4:**
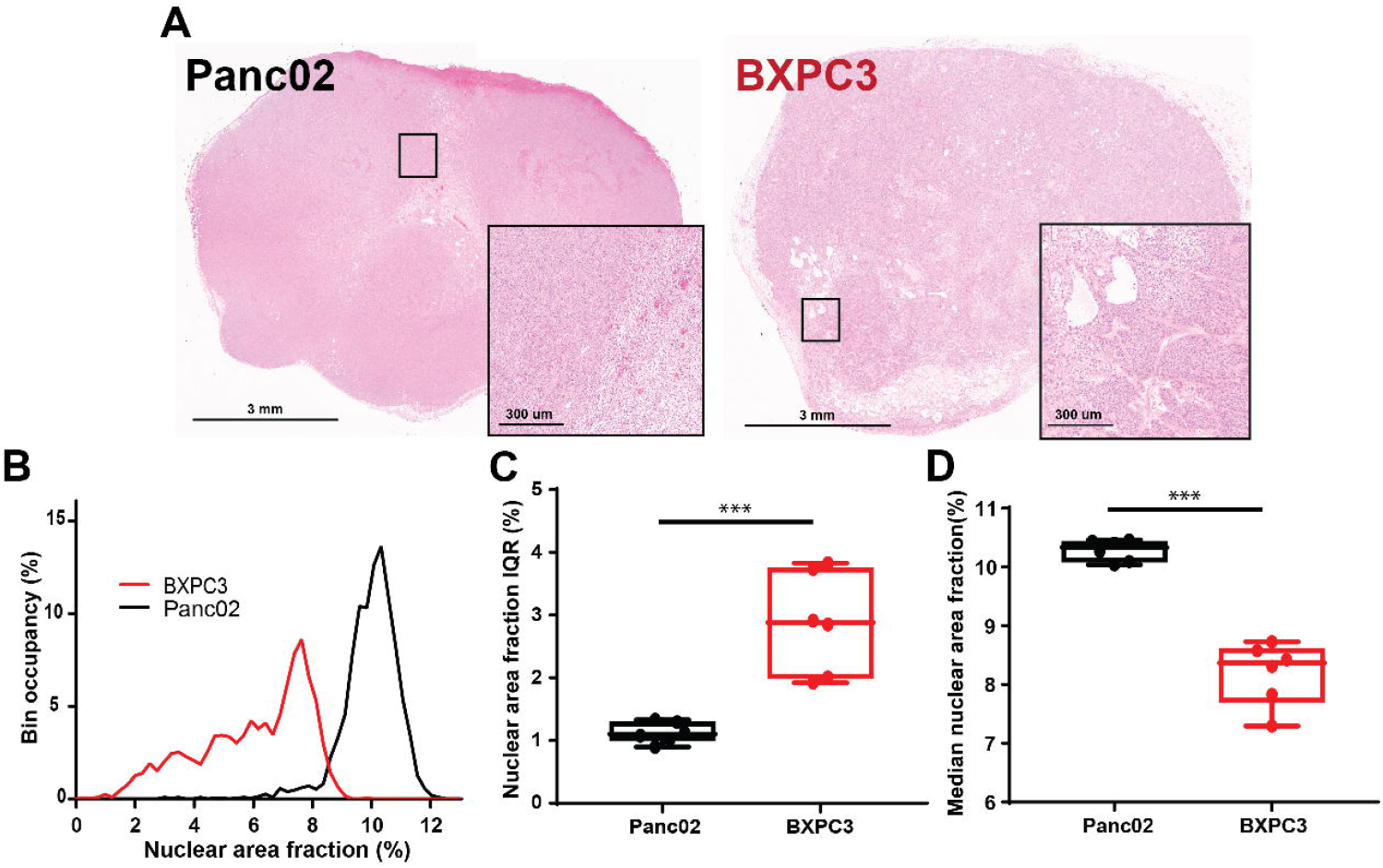
Histological comparison between two tumor types supports the imaging results. Representative H&E sections from a Panc02 (A,left) and a BXPC3 (A,right) tumors are displayed, showing a clearly less dense and more heterogeneous BXPC3 tumor. Quantification of the nuclear area fraction (as explained in Supplementary Methods) between the representative sections shows different histograms (B). Quantification of histogram metrics for representative subsets of tumors (n=6 for each type) reveals (C) a significantly a significantly higher interquartile range of the nuclear density of BXPC3 tumors, reflecting the higher heterogeneity, and a (D) lower median of the distribution. Box between 25th and 75th percentile, line at median. ***: p<0.001.

It is well established that X-irradiation causes cell death and leads to tumor necrosis (29). We postulate that the structural changes in the tumor related to local necrosis are manifested in increased heterogeneity, detected with high sensitivity by IQR quantification. To investigate this, the T2 histograms in necrotic and viable areas were compared in a subset of Panc02 tumors with well-defined necrotic regions, aided by H&E stained sections matching the MRI slices (**Figure 5A,B**). A clear difference in histograms between necrotic and viable regions was observed, as shown in a representative sample (**Figure 5C**), with the necrotic regions having a wider distribution of T2 values than corresponding viable areas. Quantification of histogram metrics confirmed these qualitative observations, a significantly higher IQR due to the elevated heterogeneity in the necrotic and peri-necrotic regions, and no clear difference in median (**Figure 5D,E**).

**Fig. 5:**
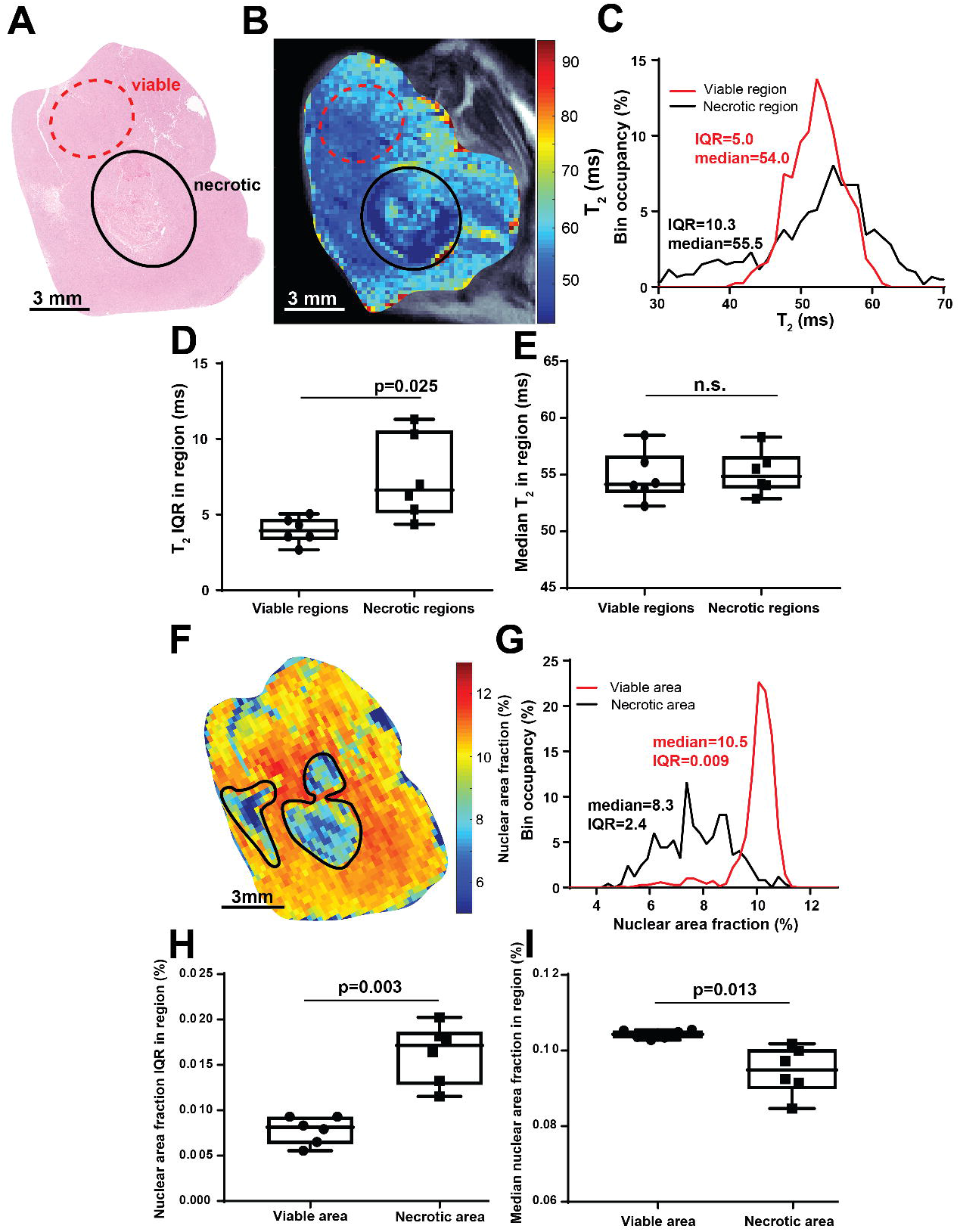
T2 histogram width is driven by tumor necrosis development. Areas of macroscopic necrosis (A, black oval) and viable tissue (A, dashed red oval) were identified in a representative H&E section, and translated onto a co-registered T2 map slice (B). The T2 histograms were calculated in these regions (C), showing different shapes. These differences were quantified for n=6 Panc02 tumors where H&E to MRI co-registration of necrotic and viable regions could be performed with minimal error. As for the representative tumor, the results show a clear difference in IQR (D) between the necrotic and viable areas, and no difference in the median T2 (E), suggesting that the (i) necrosis can be detected in T2 using histogram width instead of average intensity metrics, and (ii) the observed changes in IQR following irradiation are likely driven by necrosis development. A map of nuclear density (F, same tumor as in A-C) is shown and necrotic areas outlined in black, derived from H&E sections as described in Supplementary Methods. Corresponding value histograms in necrotic and viable areas of the section (G), show a clearly wider distribution and slight shift to lower values in necrotic regions. The histogram metrics, computed in tumors used in panels D and E, show a strongly increased IQR of the nuclear density (H) and slightly decreased median (I) in necrotic regions, in agreement with the T2 comparison from matching areas, showing that necrosis is particularly strongly manifested in high tumor heterogeneity. Box between 25th and 75th percentile, line at median.

Further histological analysis in the same tumors was directed at testing the hypothesis that the increase in T2 heterogeneity was indeed driven by the cellular heterogeneity in necrotic areas. The averaged nuclear density maps were compared in viable and necrotic regions. As is apparent from the representative image (**Figure 5F**), the necrotic areas showed generally lower values of nuclear density as well as higher heterogeneity. This observation was confirmed at the histogram level (**Figure 6G**). Similar to the previous figure, the difference in the IQR between necrotic and viable was more significant and with higher dynamic range than the median nuclear area fraction value (**Figure 6H,I**).

**Fig. 6:**
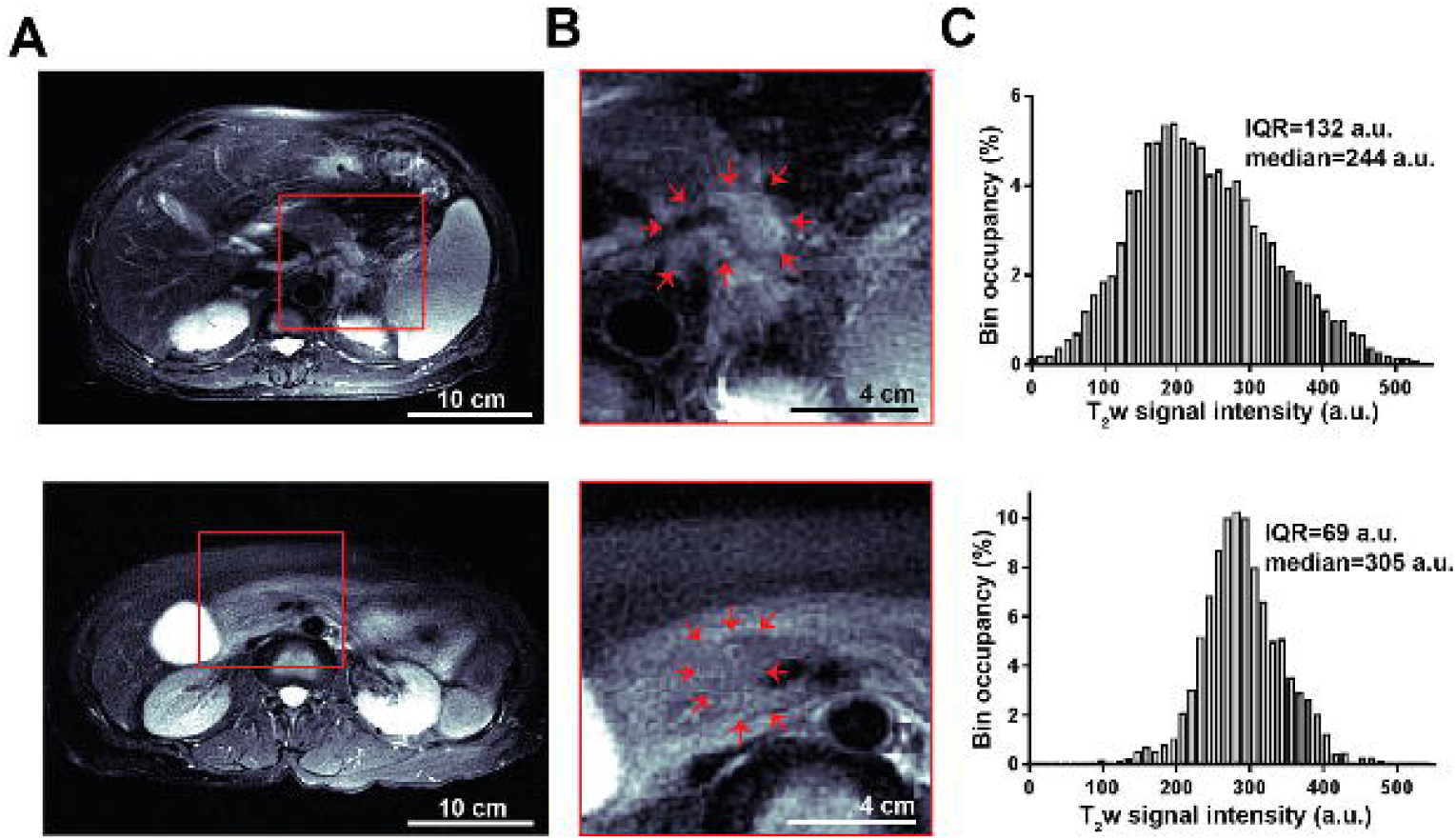
Histogram analysis enables heterogeneity quantification in clinical scans. Example axial T2 weighted images of two locally advanced pancreatic tumors (A, shown on matching intensity scales) display different level of internal heterogeneity, as seen clearly in magnified images (B), with the tumor area outlined by red arrows. The tumor in the top row shows significant heterogeneity compared to the more homogeneous mass in the bottom row, reflected in the wider distribution of signal intensity values in the histogram (C), and a corresponding higher IQR. Red box in (A) depicts the area magnified in (B).

### IQR measurements applicable in single echo T2 weighed imaging

In current clinical practice T2 images are typically acquired with a single echo time, as T2-weighted rather than quantitative T2 scans. The applicability of the above described approach was therefore assessed for raw single echo images with a TE of 77ms (**Supplementary Figure S3**). Reassuringly, in Panc02 tumors the IQR of signal intensity showed a consistent, significant increase (**Supplementary Figure S3A**) that persisted through 3 to 9 days after XRT (3.45±0.15ms vs.4.2±0.3ms (p=0.02) vs. 4.3±0.4ms (p=0.055), vs 4.2±0.3ms (p=0.01) vs 3,6,9 days respectively). The median signal intensity in Panc02 tumors showed no significant change (**Supplementary Figure S3B**) from before irradiation until 3,6 or 9 days after (p=0.98, 0.52 and 0.17 respectively). Similarly in BXPC3 the IQR increased significantly 3 days after irradiation (6.0±0.4ms vs 5.3±0.3ms, p=0.003, **Supplementary Figure S3C**), while the median showed no significant change (16.4±2.3ms vs 15.1±2.2ms, p=0.10, **Supplementary Figure S3D**). This result suggests that the IQR metric is insensitive to the signal intensity fluctuations associated with raw single echo scans. Further analysis confirmed that this conclusion applied to T2w scans acquired with a range of echo times, with IQR showing a significant change 3 days after XRT for TE=28-105ms in Panc02 tumors (for echo times 7-224ms acquired).

### IQR reflects tumor heterogeneity in clinical MR image analysis

Preliminary illustration of the clinical utility of IQR measurements for quantification of tumor heterogeneity is presented in **Figure 6**. T2 weighted MRI images of two exemplar locally advanced PDAC tumors (**Figure 6A**) show clear differences in heterogeneity. The highly heterogeneous tumor (magnified image **Figure 6B**, top row) shows corresponding wide distribution of signal values (**Figure 6C**, top row) compared to the more homogeneous tumor shown in the bottom row of **Figure 6**, reflected in the large difference in IQR (132 vs 69 a.u. top vs. bottom row).

### IQR shows excellent repeatability in vivo

Robustness of the measurements, crucial when proposing new imaging metrics, was also evaluated through test-retest measurements on a cohort of Panc02 (n=4) and BXPC3 (n=4) tumors. These animals were removed from the MRI after imaging, repositioned in the holder, and scanned again. The relationship between the image features hence measured is a good estimate of the feature robustness (30). Overall a good test-retest correlation between the measurements was observed for both IQR and median (Pearson r=0.998 and r=0.949 respectively, **Supplementary Figure S4**). Interclass correlation coefficient, commonly used for test-retest comparison (28) confirmed the excellent robustness of the IQR, superior to the median and other histogram metrics (0.997 vs. 0.896, 0.907, 0.867, 0.207, 0.198 interclass coefficient (ICC) for IQR vs. median, mean, standard deviation, kurtosis and skewness respectively), suggesting again that IQR may be a suitable imaging biomarker candidate in tumor studies. Despite the use of arbitrary units in place of absolute relaxation times, the use of single echo T2-weighted scans only slightly reduced the test-retest agreement of IQR (ICC 0.858).

## Discussion

The main goal of the current study was to identify and develop a clinically-translatable metric of tumor structural heterogeneity that is highly robust and sensitive for detection of changes in cellularity and viability of the tumor early after irradiation.

T2-weighted imaging is widely used clinically for delineation of anatomical structures and detection of pathological changes, both in cancer and in other diseases. Fast, robust sequences such as the Fast Spin Echo (31) are well established in the field, yet quantitative analysis of the T2w images is not practiced, in contrast to, e.g. ADC maps. In this manuscript we hypothesized that T2 relaxation contrast, in conjunction with a novel analysis method focused on structural heterogeneity, may be highly sensitive and reproducible for measurement of early radiotherapy-induced changes. Interquartile range (IQR) of T2 is presented as a promising imaging biomarker candidate of tumor heterogeneity, informative of radiation response, and its technical and biological validation was performed, as is needed for all biomarkers (32). The metric is contrasted with other histogram features, and in particular the median tumor signal, which provides the whole-tumor insight, reinforcing the importance of heterogeneity quantification provided by the IQR.

Imaging of tumor heterogeneity is an emerging approach in the field of imaging, aimed to maximize the information extracted from radiological data by expanding the analysis from traditional average tumor values and hotspot detection to take into account the spatial and value distribution of voxels (3). The results presented in this work support this view. We found that quantification of the interquartile range (IQR) of T2 values, reflecting the intra-tumor heterogeneity, enabled powerful insight into the radiation-induced changes in cellularity and viability, with sensitivity superior to the commonly considered first order metrics. In the murine models considered, the measured cellular heterogeneity changes were related to tumor volume response to radiation. In other cancers, early cellularity modulation after radiation therapy, as measured by Diffusion Weighted MRI, were indicative of clinical response (33–35). The presented data suggests that T2 IQR may offer a more sensitive measure of these changes, which in the future can be exploited for temporal adaptation of the radiotherapy dosing and fractionation scheme. This approach, already used with promising results in chemotherapy (36,37), could provide a paradigm shift in radiotherapy planning, offering personalization of XRT schedule for long term disease control minimized doses. Future work will focus on application of the findings for this clinical unmet need (38).

The choice of 10 Gy radiation dose in the main experiment presented was made to match the dose per fraction of the clinical MR-Linac pancreatic cancer treatment protocol as often used at and tested in clinical trials (39). The rapidly increasing use of MR-Linac systems, offering MR imaging insight into the tumor at each fraction, creates an opportunity and an unmet clinical need for quantitative analysis tools enabling detection and of the tumor radiotherapy response correlated to later tumor growth. The approach presented in this study, especially given its sensitivity, robustness, and applicability to single echo images, has the potential to allow us to personalize radiotherapy treatment.

Quantitative histological image analysis was used to shed light on the informational content of T2 IQR and validate its biological underpinnings, a challenging task in radiomic and image feature extraction studies. Necrotic areas showed a drastically wider value distribution than viable regions, both in T2 maps and in *ex vivo* cellularity measurements, while remaining closely matched in terms of average values. These findings reinforce the understanding that the strong changes in T2 IQR following irradiation were caused by early necrosis formation, characterized by a significantly increased structural heterogeneity, which can be detected with higher sensitivity than the corresponding changes in mean values. In addition, comparing the two tumor types in terms of T2 and histological metrics, we showed quantitative differences consistent with our understanding of IQR as a measure of structural heterogeneity at the spatial scale of the MR voxel size. The highly homogeneous Panc02 tumors as seen from H&E staining show low T2 IQR and its strong increase after irradiation, compared to the more differentiated BXPC3 tumors, with pockets of fluid and necrosis. Proof-of-concept analysis of clinical images confirmed that differences in IQR are consistent with visual tumor heterogeneity.

In current clinical practice, quantitative T2 relaxation measurements are rarely performed in cancer imaging. Instead, T2-weighted scanning is used for qualitative assessment of the soft tissue structure. This approach however suffers from lack of internal normalization, with signal intensity depending on the electronic gain and pulse power. The use of IQR as shown in the study may overcome this limitation, as it showed robustness and sensitivity to radiotherapy related changes in single echo T2 weighted images, with no need for normalization. This observation is crucial for the field of quantitative MRI and MRI radiomics, as signal normalization remains one of the main challenges for meaningful feature extraction and comparison (40). In addition, it highlights the readily translatable nature of the method described. Going forward we postulate however, that quantitative T2 relaxation measurements, as often acquired in cardiac MRI (41) and shown in prostate cancer (42) as part of multiparametric MRI (43), could be performed routinely in the clinic, in combination with heterogeneity-focused analysis. As shown in this study, absolute T2 relaxation measurement, and associated IQR quantification, can provide highly robust and sensitive measure of the radiation induced viability and structure changes, exceeding the ability of ADC, routinely used for quantification of macroscopic necrosis.

There are some limitations to the presented work. T2 contrast, although widely used in cancer imaging, is less well defined in terms of its biological meaning than diffusion contrast for assessment of tissue structure, as it can be affected by multiple other factors such as swelling or presence of blood and fat. Further measurements may be required to quantify the influence of these elements in the context of radiation response. The experiments were performed in subcutaneous model which does not fully recapitulate the clinical disease. Immunocompetent and deficient models were utilized to include the confounding effects of immune system, yet different pattern of immune infiltration and inflammation following irradiation in humans compared to mice may alter the dynamics of imaging changes. Validation in a more realistic transgenic model (44), as well as in retrospective clinical MR-Linac scans are planned in the near future.

In summary, the study presents a novel, translatable method and an associated imaging biomarker for measurement of early radiation induced changes in tumor cellularity and viability in murine models of pancreatic cancer, relevant for treatment response. The study highlights the value of sophisticated histogram analysis and heterogeneity quantification for extraction of biologically relevant information from a standard and widely used Fast Spin Echo MRI sequence. We envision the findings to be swiftly applied to clinical data, in particular with the advent of the MR-Linac systems, acquiring images as often as daily with each radiation fraction.

## Acknowledgements

This work has been supported in part by the Analytic Microscopy Core Facility, Small Animal Imaging Laboratory Core Facility and Tissue Core Facility at the H. Lee Moffitt Cancer Center & Research Institute; an NCI designated Comprehensive Cancer Center (P30-CA076292).

## Notes

**Funding:** This work was supported by NIH grants U01 CA143062, U54 CA143970, core facilities supported by the CCSG core grant P30 CA76292

Dr. Gillies serves on the Advisory Board for HealthMyne Inc. Other authors declare no potential conflicts of interest.

https://github.com/mrtomasz91/T2IQRData

### Competing Interest Statement

Dr Gillies is an investor and serves on an advisory board for HealthMyne Inc.

