## Supplemental Materials for "T2 heterogeneity provides a sensitive measure of early tumor response to radiotherapy"

**Supplementary Materials:**

1. **Supplementary Figures**
2. **Supplementary Tables**
3. **Supplementary Methods**

**Supplementary Figures**

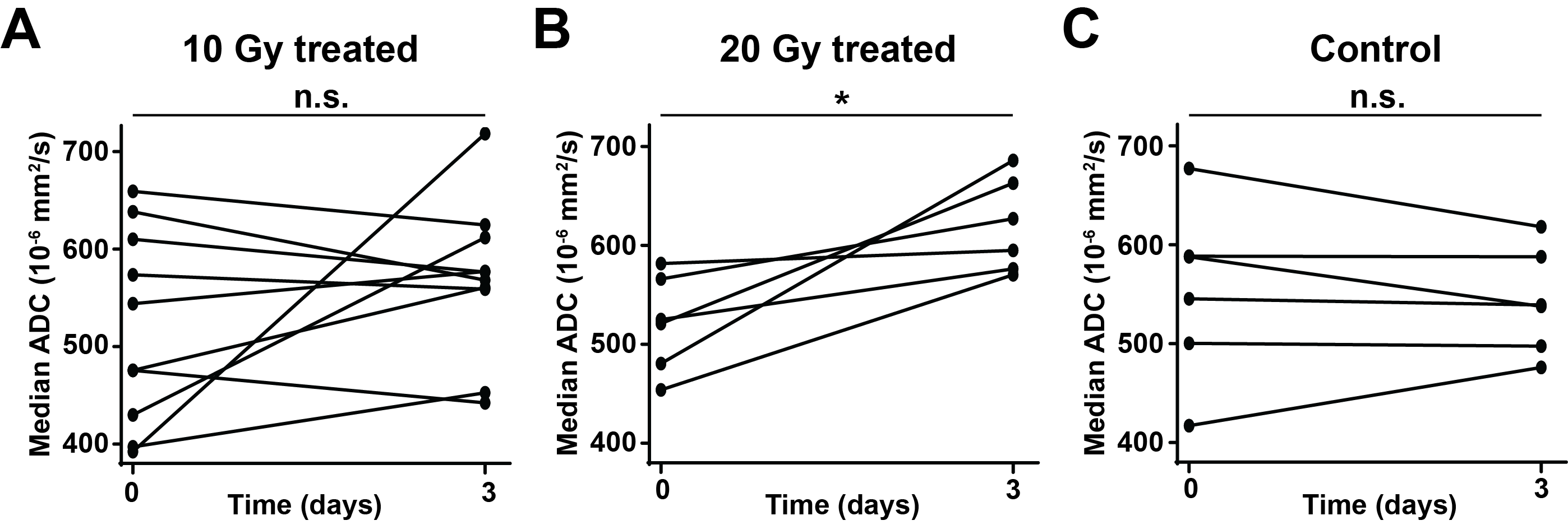

Supplementary Figure S1: **Diffusion Weighted MRI shows modest changes in response to radiotherapy.** Median Apparent Diffusion Coefficient (ADC) in the tumor showed no significant change 3 days after 10 Gy irradiation (A). Only increasing the dose to 20 Gy in a separate cohort lead to a significant median ADC increase (B). No changes were observed in control, sham irradiated tumors (C). n=11, 6 and 6 Panc02 tumors for 10 Gy, 20 Gy and control respectively. n.s: non-significant, *:p<0.05.

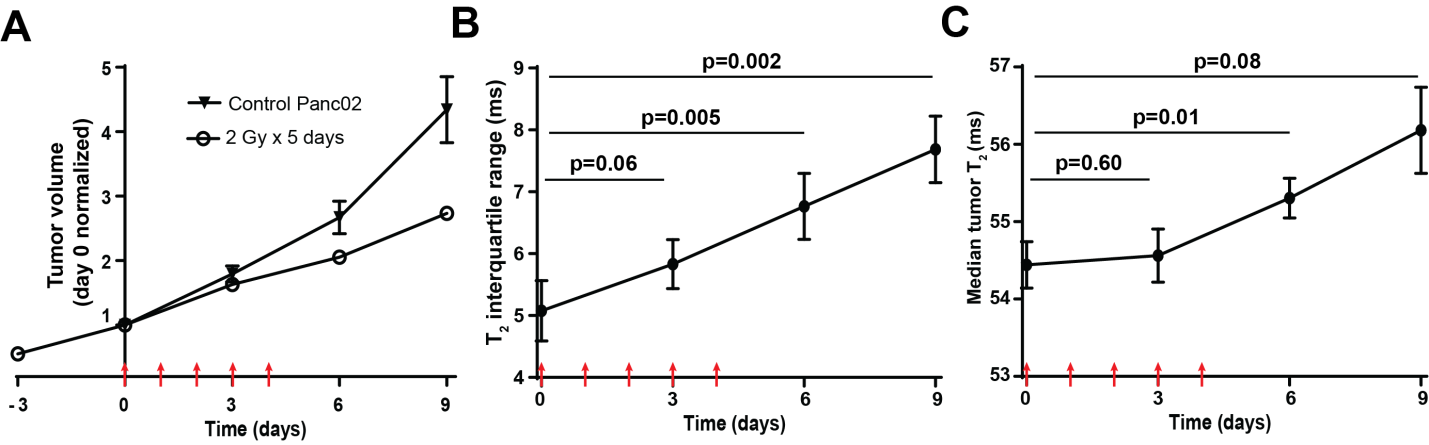

Supplementary Figure S2: **Traditional fractionation supports single dose results.** Treatment with 5 daily fractions of 2 Gy lead to a moderate slowdown of tumor volume increase (A) in Panc02 tumors. The T2 interquartile range (B) displayed clear increase as early as 3 days after treatment start, while the median (C) shows modest, delayed changes. P values calculated by two-sided paired t-test compared to day 0. Tumor volume normalized to day 0 in treated (n=6) and control (n=6) to highlight growth trends. Red arrows indicate irradiation (2Gy). Error bars denote standard error of the mean, with lack of thereof indicates error smaller than point size.

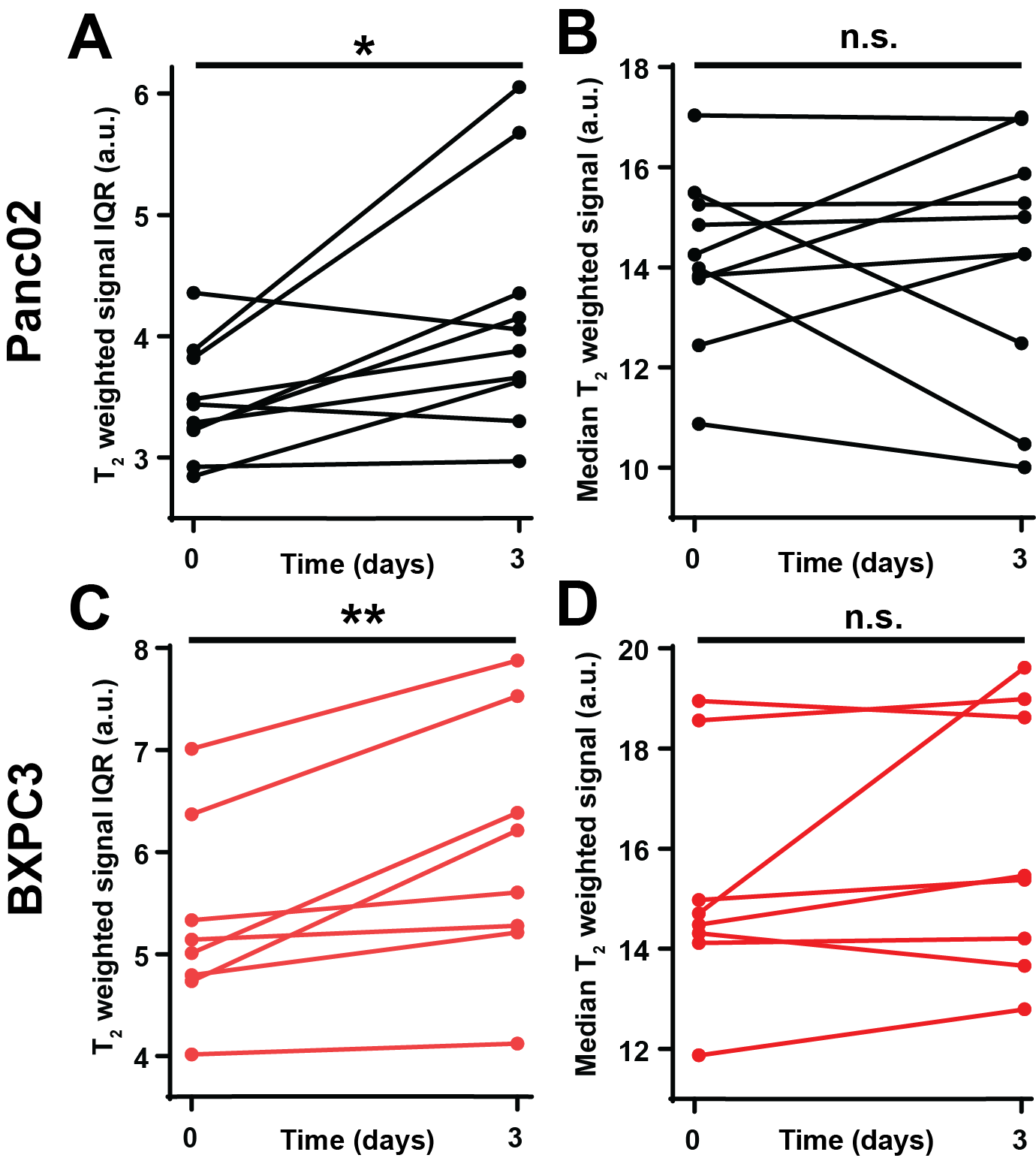

Supplementary Figure S3: **T2 weighted scans show changes in IQR after irradiation.** 3 days post radiotherapy, tumor signal IQR from T2 weighted scans (A) inPanc02 tumors showed significant increase matching this of T2 maps, while median signal (B) showed no change. Similar results were seen in the IQR (C) and median (D) in BXPC3 tumors. n.s. denotes not statistically significant, * p<0.05, **p<0.01.

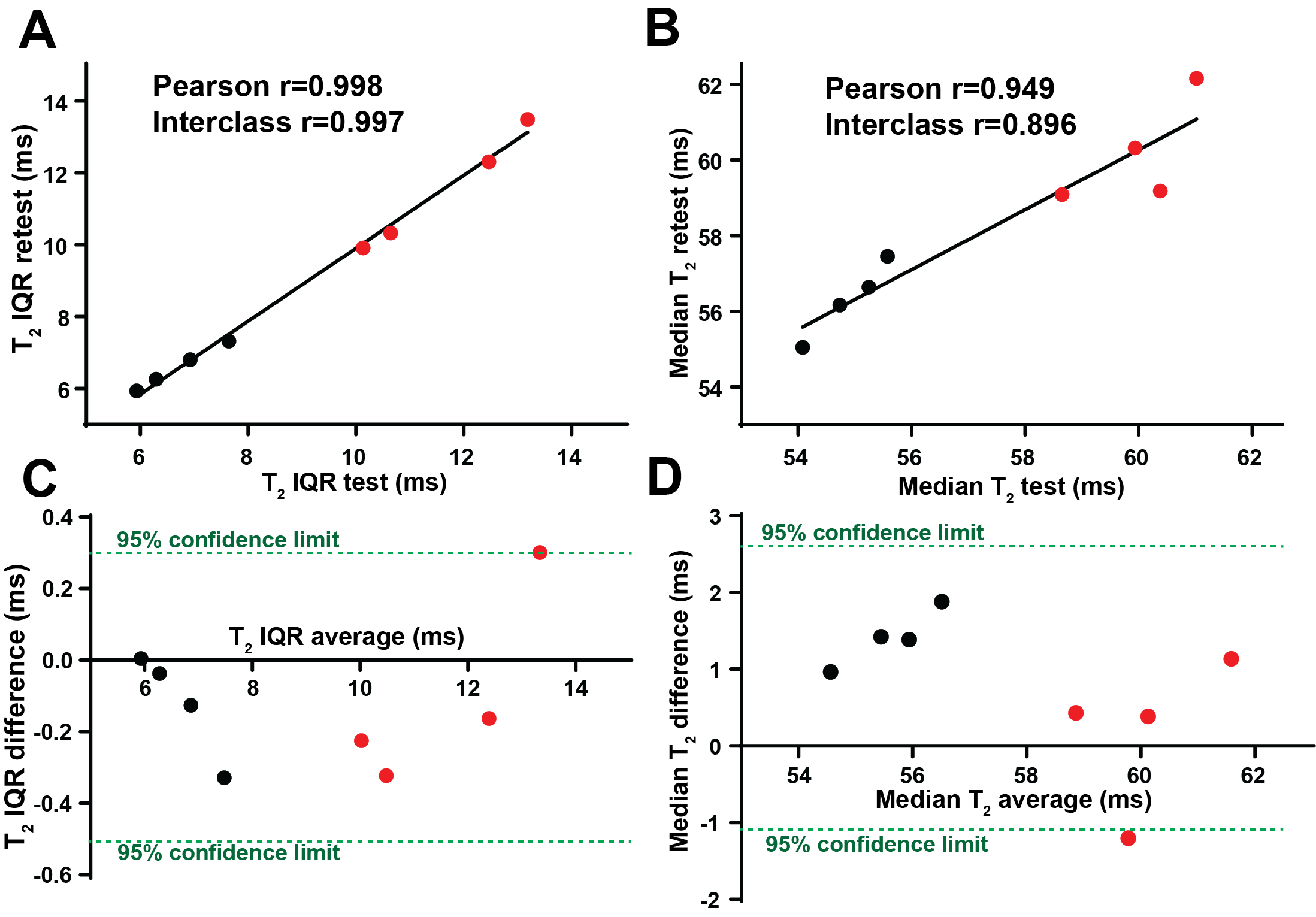

Supplementary Figure S4: **T2 IQR is a highly robust image feature**. Results of test-retest measurements in a cohort of n=4 Panc02 (black points) and n=4 BXPC3 (red points) were quantified and plotted. Excellent correlation for T2 IQR (A) exceeded this of median T2 (B), a standard whole-tumor metric, suggesting superior reliability of the novel IQR metric. The Bland-Altman plots for median (C) and IQR (D) are also shown, presenting a good test-retest agreement of both metrics, mostly fitting within the 95% confidence limits shown in green dotted line.

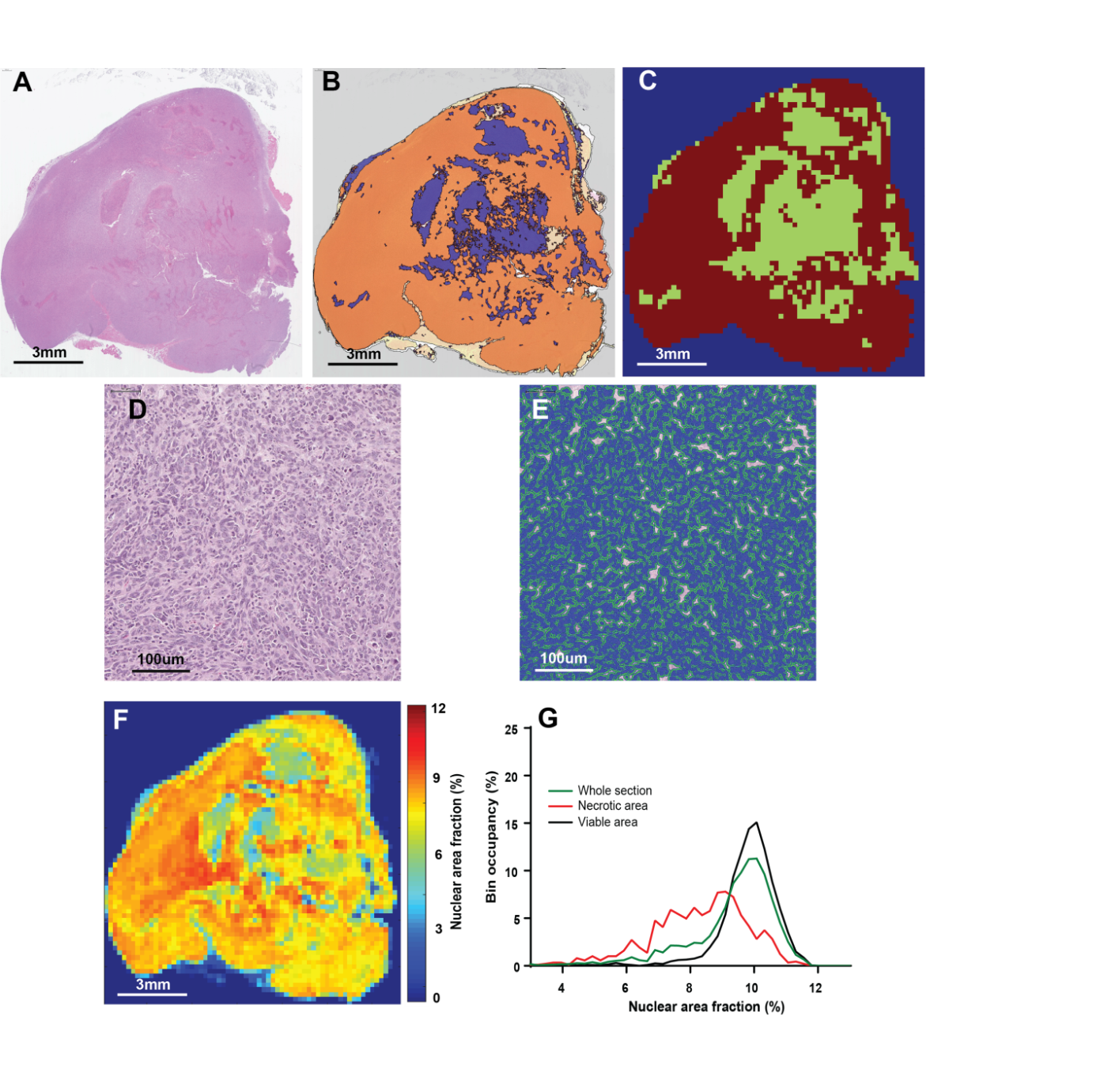

Supplementary Figure S5: **Quantification method for cell density from H&E sections**. A H&E stained section was analyzed in Tissue Studio software to automatically identify necrotic (B, blue mask) and viable (B, orange mask) areas. The machine learning algorithm was trained on small patches of tissue manually identified by an experienced user. For further analysis the high resolution masks were loaded into MATLAB, and down-sampled to match the resolution of T2 maps in MRI (C). Next, all cells in the section (D) were identified and segment automatically in Tissue Studio (E) into nuclei (blue), cytoplasm (green) and other non-cell areas (white). The high-dimensional list of nuclei positions and areas was loaded into Matlab, where a low-resolution image of pixel area fraction taken up by nuclei (nuclear area fraction) was calculated (F). This image could then be used as a proxy for cellular density in the region. Given the necrotic/viable segmentation, histograms of nuclear area fractions in each region were calculated for each section (G).

**Supplementary Tables**

| **Panc02 tumors**  **(10 Gy XRT)** | **p value** | **Value pre XRT (ms)** | **Value 3d post XRT (ms)** | **Ratio 3d/pre XRT** |
| --- | --- | --- | --- | --- |
| **Interquartile Range** | **0.001** | **5.4±0.3** | **6.9±0.5** | **1.27±0.06** |
| **Median** | **0.01** | **54.6±0.5** | **55.8±0.6** | **1.025±0.007** |
| **Mean** | **0.04** | **56.0±0.7** | **58.3±0.9** | **1.05±0.02** |
| Kurtosis | 0.21 | 40±14 | 34±3 | 0.91±0.10 |
| Skewness | 0.89 | 4.0±0.4 | 4.0±0.3 | 1.01±0.08 |
| Standard Deviation | 0.12 | 9.3±0.8 | 12.6±2.1 | 1.6±0.3 |

**Supplementary Table 1:** Summary of histogram feature values in 10 Gy treated Panc02 tumors (n=11) before and 3 days after irradiation. P value by paired t-test, standard error of the mean quoted. Values in bold denote significant changes.

| **Panc02 tumors (control)** | **p value** | **Value pre XRT (ms)** | **Value 3d post XRT (ms)** | **Ratio 3d/pre XRT** |
| --- | --- | --- | --- | --- |
| Interquartile Range | 0.55 | 5.3±0.3 | 5.5±0.2 | 1.05±0.07 |
| Median | 0.71 | 56.1±0.8 | 55.7±0.4 | 0.994±0.017 |
| Mean | 0.41 | 58.1±0.8 | 57.1±0.5 | 0.984±0.018 |
| Kurtosis | 0.34 | 33±3 | 68±4 | 2.0±0.9 |
| Skewness | 0.47 | 4.5±0.4 | 5.5±1.4 | 1.2±0.3 |
| Standard Deviation | 0.81 | 19±3 | 18.7±1.2 | 0.99±0.17 |

**Supplementary Table 2:** Summary of histogram feature values in sham treated Panc02 tumors (n=6) before and 3 days after sham irradiation. P value by paired t-test, standard error of the mean quoted.

| **BXPC3 tumors**  **(10 Gy XRT)** | **p value** | **Value pre XRT (ms)** | **Value 3d post XRT (ms)** | **Ratio 3d/pre XRT** |
| --- | --- | --- | --- | --- |
| **Interquartile Range** | **0.01** | **12.6±0.6** | **14.2±0.8** | **1.13±0.04** |
| Median | 0.08 | 58.4±0.7 | 59.9±0.7 | 1.025±0.012 |
| Mean | 0.46 | 63.8±1.4 | 65.1±1.0 | 1.02±0.02 |
| Kurtosis | 0.76 | 19±3 | 20±4 | 1.3±0.3 |
| Skewness | 0.84 | 3.2±0.3 | 3.2±0.3 | 1.06±0.13 |
| Standard Deviation | 0.83 | 19±3 | 18.7±1.2 | 1.07±0.11 |

**Supplementary Table 3:** Summary of histogram feature values in 10 Gy treated BXPC3 tumors (n=8) before and 3 days after irradiation. P value by paired t-test, standard error of the mean quoted. Values in bold denote significant changes.

| **BXPC3 tumors**  **(control)** | **p value** | **Value pre XRT (ms)** | **Value 3d post XRT (ms)** | **Ratio 3d/pre XRT** |
| --- | --- | --- | --- | --- |
| Interquartile Range | 0.63 | 16±3 | 14±2 | 0.94±0.17 |
| Median | 0.86 | 57.6±0.8 | 57.5±0.9 | 0.997±0.01 |
| Mean | 0.75 | 66±3 | 64±2 | 0.98±0.06 |
| Kurtosis | 0.29 | 15±3 | 22±6 | 1.6±0.4 |
| Skewness | 0.24 | 2.8±0.2 | 3.3±0.3 | 1.21±0.14 |
| Standard Deviation | 0.89 | 24±7 | 23±4 | 1.1±0.3 |

**Supplementary Table 4:** Summary of histogram feature values in sham treated BXPC3 tumors (n=4) before and 3 days after sham irradiation. P value by paired t-test, standard error of the mean quoted.

| **Slide Scanner** | Leica Aperio™ AT2 (Vista, CA) |
| --- | --- |
| **Objective** | 20x/0.8 NA Plan Apo |
| **Resolution** | 0.5022 µm/pixel |
| **Analysis Software** | Definiens Tissue Studio v2.7 (Munich, Germany) |
| **Analysis Magnification** | 20 |
| **Algorithm** | Nucleus Detection |
| **Hematoxylin Threshold** | 0.05 |
| **Typical Nucleus Size** | 40 µm² |
| **Nuclear Size Threshold** | Area > 10 µm² |

**Supplementary Table 5:** Parameters used for H&E quantification.

**Supplementary Methods**

**Histological analysis**

H&E stained sections were segmented and analyzed in Tissue Studio v2.7 (Definiens, Munich, Germany) software to quantify local nuclear density distribution. Nuclei were identified based on hematoxylin staining, and their position and area recorded (see **Supplementary Table 5** for parameters). A built-in machine learning algorithm was also used to detect necrotic and viable areas of the tissue, based on manual classification of small areas of the sections. The list of nuclei and viability map were then loaded into MATLAB 2018b (Mathworks, Natick, MA), where a map of voxel area occupied by nuclei was computed, in resolution matching that of T2 maps, to provide histological estimate of the local cellular densities, as observed in MRI. A co-registered necrosis mask allowed comparison of the image value statistics and histograms in viable and necrotic areas of each section. The procedure is further explained in **Supplementary Figure S5**.

**Magnetic Resonance Imaging**

For MR imaging, mice were anaesthetized with isoflurane (3% for induction, 2-2.7% for maintenance, mixed in pure oxygen), and placed into the bore of the magnet (Bruker, 7T) in supine position, with the tumor area placed in the middle of the coil (Whole body quadrature coil, 35mm diameter, M2M). Following anatomical coronal and axial scans (T2 weighted, FSE, TR/TE=1800/30 ms, 256x256 points, 35mm FOV, 1/0 mm slice thickness/gap), T2 map was acquired of the entire tumor volume (MSME, TR=3427ms, 32xTE=7-224ms, 7ms spacing, 35 mm field of view,128x128 points, 1/0 mm slice thickness/gap). Diffusion Weighted Imaging was performed on a sub-cohort of the animals using an Echo Planar Imaging sequence (TR/TE=1600/42ms, 3 directions, b- values 100,450,700 s/mm^2^, respiratory gating).
